# Inactivated SARS-CoV-2 Reprograms the Tumor Immune Microenvironment and Improves Murine Cancer Outcomes

**DOI:** 10.1101/2022.06.30.498305

**Authors:** Eileena F Giurini, Michael Williams, Adam Morin, Andrew Zloza, Kajal H Gupta

**Author notes:** Contributed equally (Eileena F Giurini, Michael Williams).

## Abstract

Following the breakthrough of immune check point inhibitors (ICIs), a new era of immuno-oncology agents has emerged and established immunotherapy as a part of cancer treatment. Despite the improving outcomes of ICIs, many patients with initial response are known to develop acquired resistance later. There is increasing interest in utilizing other stimulatory means, such as anti-pathogen immune responses to induce anti-tumor immune responses. The immunostimulatory effects of anti-pathogen-treated tumors in combinations with ICI are known to potentially amplify anti-tumor immunity resulting in increased tumor responses and improved outcomes. Anti-pathogen-treated tumors can become immune-infiltrated “hot” tumors and demonstrate higher treatment response rates and improved survival. Our research group has previously demonstrated that tumors can be converted from “cold” to “hot” by intratumoral injection of a commercially available seasonal influenza vaccine. In continuation with our work, in deciphering the role of anti-viral immunity in the context of tumor immunology, we studied the role of inactivated SARS-CoV-2 virus as anti-tumor agent. Here we report that intratumoral injections of inactivated SARS-CoV-2 convert the immunologically cold tumors to hot by generating anti-tumor-mediated CD8^+^ T cells. Our findings suggest that inactivated SARS-CoV-2 can be used as an immune modulator in immunotherapy for melanoma and triple-negative breast cancer.

## Introduction

The last decade has experienced a major addition to the cancer therapy arsenal. Antibody-based immunotherapies have been introduced to exploit the precision of the immune system to specifically target tumor cells and cancer-driving immune cell populations. Thus, immune checkpoint inhibitors (ICIs) have rapidly altered the treatment topography of multiple tumor types. Despite the characteristic durability of response to ICI, significant variability in effectiveness still exists between cancers and even between individual patients with the same cancer type.

The heterogeneity in response to ICIs is due to multiple factors, which may be related to differences in the composition of the tumor immune microenvironment (TIME). TIMEs can be classified broadly into two categories: those that are devoid of immune cells or those that are immune cell-infiltrated. Further, the infiltrated TIMEs may contain different cell populations; some may be inflamed with pro-immune cytotoxic lymphocytes, while others may be saturated with immunosuppressive cell populations. Previous studies have demonstrated that different TIMEs may result in different effects of immunotherapy. Checkpoint inhibitors produce greater anti-tumor effects when the TIME is “hot,” or inflamed with pro-immune cell, compared to “cold” tumors that are less immunogenic.

In December 2019, our group published a study which demonstrated that the seasonal influenza vaccine can be repurposed as a cancer immune therapy^1^, where intratumoral administration of the vaccine converted immunologically “cold” tumors to “hot.” Host response to influenza virus enhanced antitumor immunity, notably thorough an increase in tumor-specific CD8+ T cells. Additionally, influenza vaccine treatment sensitized tumors to immune checkpoint inhibitor therapy. Notably, these effects were only seen after injection with the non-adjuvanted vaccine containing heat-inactivated influenza virus, suggesting that some vaccines may be more immunogenic than others.

Our study on intratumoral injection of heat-inactivated influenza virus as an immunotherapy agent was immediately followed by the onset of the COVID-19 pandemic. The rapid spread of SARS-CoV-2 established itself as a highly immunogenic virus, with notable similarities to influenza, including both viruses are single-stranded RNA viruseses^2^. Both viruses infect the respiratory tract and use surface proteins to infect the host. Influenza requires hemagglutinin^3^ and neuraminidase for host cell invasion, whereas SARS-CoV-2 relies on protein S. Both viruses depend on a viral RNA polymerase to express their proteins. Given that several clinical cases have been reported where in COVID-19 infections were associated with tumor regression in melanoma^4^ and Hodgkin Lymphoma^5, 6^, we hypothesized that a host response to intratumoral injection of chemically inactivated SARS-CoV-2 may produce an anti-tumor effect in mouse models akin to what we observed with influenza. In this study our findings propose that inactivated SARS-CoV-2 vaccines may have the potential for use as a cancer immunotherapy.

## Materials and Methods

### Mice

BALB/C and B6 (C57BL/6J) mice were purchased from The Jackson Laboratory at 6-8 weeks of age. All animals were housed in specific pathogen-free facilities and all experimental procedures were in accordance with policies approved by the Institutional Animal Care and Use Committee (IACUC) and Rush University Medical Center.

### Inactivated SARS-CoV-2 virus

Chemically inactivated COVID-19 was purchased from Microbiologics San Diego. For TLR-activation assays, 1.5 × 10^6^ plaque-forming units (PFU) per mL of inactivated COVID-19 was used. For all animal experiments, mice were administered 50 µl of inactivated COVID at a concentration of 1 × 10^6^ PFU/mL via intratumoral injection. Control mice received the same volume of PBS via the same route.

### Seasonal Influenza Vaccine

Fluarix Quadrivalent (GlaxoSmithKline), a 2020-2021 FDA-approved seasonal influenza vaccine was purchased for these studies: 50 µl of influenza vaccine or a PBS control was administered to mice via intratumoral injection. For TLR activation assays, Fluarix was diluted 1:2 in PBS prior to plating.

### Cell Culture

HEK-Blue TLR reporter cell lines (Invivogen) were transfected with plasmids containing a murine TLR and an NF-κB inducible secreted embryonic alkaline phosphatase (SEAP) reporter gene, used to determine TLR stimulation. mTLR7, and non-TLR expressing parental cell line, Null2k, (Invivogen) were cultured using DMEM (Gibco), supplemented with 10% Fetal Bovine Serum (Corning), 100 units/mL penicillin (Gibco), 100 mg/mL streptomycin (Gibco), and 100 µg/mL normocin (Invivogen). Expression of TLR and SEAP plasmids was maintained with the addition of selective antibiotics, blasticidin (30 µg/mL, Invivogen) and zeocin (100 µg/mL, Invivogen). The murine triple-negative breast cancer cell line, 4T1, was cultured in RPMI (Gibco), supplemented with 10% Fetal Bovine Serum, 100 units/mL penicillin (Gibco), 100 mg/mL streptomycin (Gibco), and 100 µg/mL normocin (Invivogen). The murine melanoma cell line B16-F10 was cultured in DMEM supplemented with 10% FBS, 100 units/mL penicillin, 100 mg/mL streptomycin, and 100 µg/mL normocin.

### TLR Activation Assay

To evaluate TLR7 stimulation, 20 µl of inactivated COVID-19 or influenza vaccine was plated in duplicate or triplicate in 96-well plates. TLR7 agonist (CL264, 50 μg/mL) was run simultaneously as a positive control. 1X PBS was used as a negative control. After plating the treatments, 50,000 mTLR7 or Null2k cells suspended in 180 µl HEK-Blue Detection (Invivogen) medium were added to the respective wells. All cells were incubated for 24 hours at 37°C and 5% CO_2_. Following incubation, TLR stimulation was quantified using a Cytation 3 plate reader (BioTek) measuring absorbance at 620 nm.

### Tumor Challenge

BALB/C mice were anesthetized with isoflurane and injected with 500,000 4T1 triple-negative breast cancer cells in 100 µl PBS in the mammary fat pad (MFP). B6 mice were anesthetized with isoflurane and challenged with 100,000 B16-F10 melanoma cells in 100 µl PBS via intradermal injection. Tumor development was monitored using Vernier calipers, where tumor area was determined by two measurements in perpendicular directions. To comply with IACUC policies, tumor-bearing mice were humanely sacrificed upon tumor measurements reaching 15 mm in either direction.

### Flow Cytometry

Tumor-bearing mice were humanely sacrificed via carbon dioxide inhalation. Tissues were processed as previously described^1^. Extracellular staining for flow cytometry was performed with antibodies targeting CD3, CD4, CD8, CD11b, CD11c, CD20, CD45, Ly-6G/Gr-1, and MHC-II. All antibodies were purchased from BioLegend, BD, eBioscience, or R&D Systems. An extracellular stain cocktail comprised 1-5 µl/test combined with 0.25 µl/test of Live/Dead Fixable Aqua Dead Cell Stain Kit (405 nm excitation) (Invitrogen) in a total volume of 100 µl (made in PBS). The extracellular stain cocktail was added to each sample and subsequently incubated in a dark environment at room temperature for 30 minutes. Following extracellular staining, samples were washed twice with 200 µl PBS. For optimal intracellular staining, samples were permeabilized and fixed with 100 µl of Cytofix/Cytoperm^1^ for 15 minutes at 4°C. Samples were subsequently washed with 100 µl of 1X Perm/Wash Buffer ^1^. Samples were incubated with intracellular stain for 30 minutes at 4°C, protected from light. Samples were subsequently washed with 200 µl of 1X Perm/Wash Buffer twice, followed by two 200-µl PBS washes. Flow cytometry was performed on the BD LSRFortessa. Flow cytometry analysis was completed using FlowJo (version 10).

### Statistical Analysis

Statistical analysis was performed using GraphPad Prism (version 9.1.2). For studies involving more than two groups, a 1-way ANOVA with Tukey correction was used to determine statistical significance. A two-way ANOVA was performed to determine statistical significance with Sidak’s correction for studies with multiple time points.

## Results and Discussion

### Intratumoral injection of inactivated SARS-CoV-2 reduces tumor growth in murine melanoma and triple negative breast cancer models

We previously reported that the growth of B16 melanoma tumors could be halted by intratumoral injection of an inactivated influenza vaccine^1^. Using the B16-F10 murine model, we compared the growth patterns of melanoma tumors after injecting either inactivated influenza [encompassed within a seasonal influenza vaccine (FluVx)], inactivated SARS-CoV-2 (iCOVID), and buffer controls. SARS-CoV-2 caused tumor growth inhibition similar to that of influenza, both of which significantly halted growth compared to controls (Figure 1A). We then asked whether the effects of inactivated viruses were unique to melanoma, or if tumor growth may be affected in other cancer types. We elected to study triple-negative breast cancer, as breast cancer would be ideal for clinical translation given that tumors are typically superficial and could be easily accessed for intratumoral treatment in patients. After establishing tumors in mice using the 4T1 triple-negative breast cancer (TNBC) model, we repeated the same protocol by injecting FluVx, iCOVID, and negative control. FluVx and iCOVID were both observed to dramatically reduce tumor growth compared to control; tumors in the FluVx or iCOVID treated groups were significantly reduced of those in the control group (Figure 1B). These findings suggest that introduction of iCOVID in the tumor can elicit antitumor responses similar to that of FluVx.

**Figure 1:**
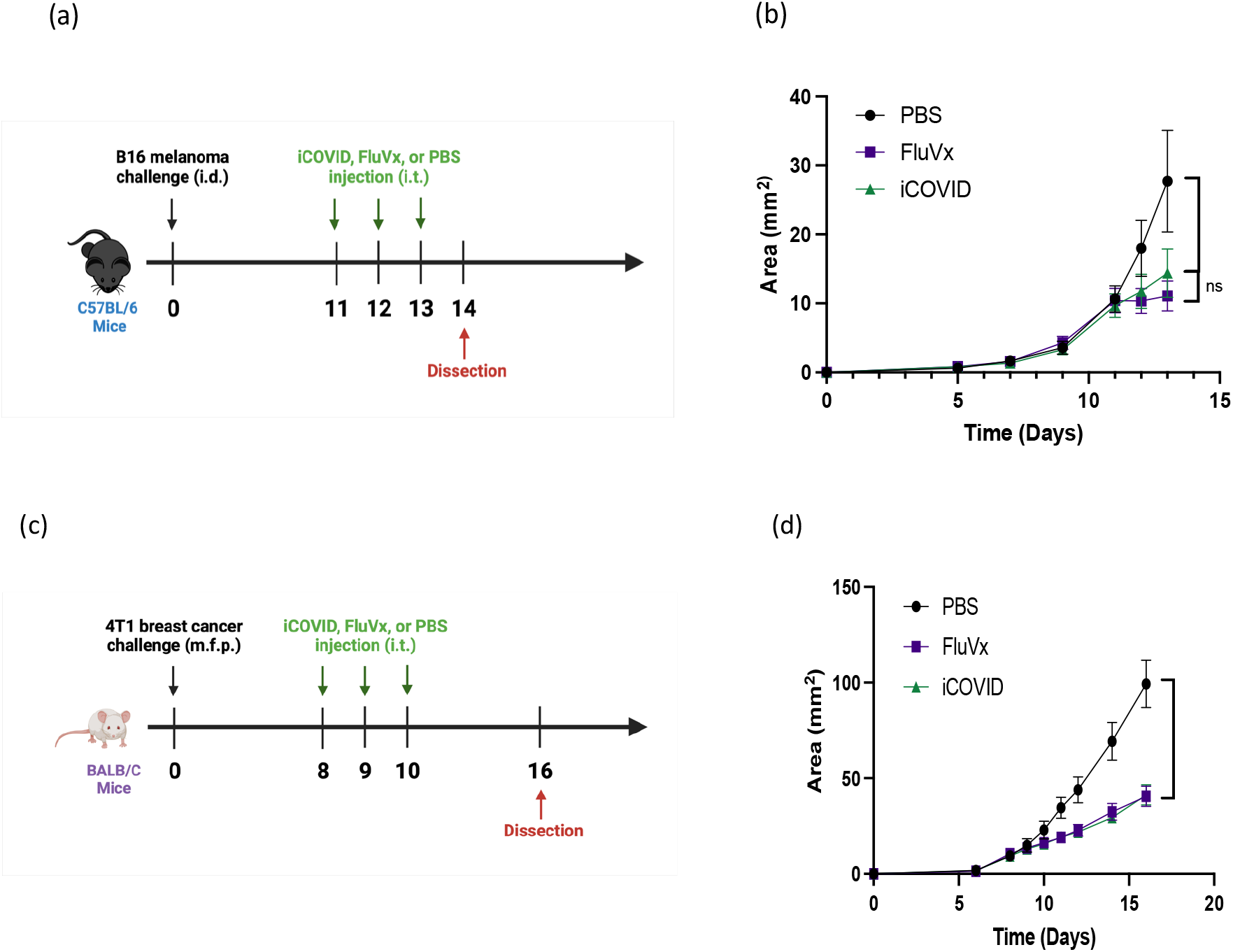
Intratumoral administration of inactivated SARS-CoV-2 reduces tumor growth. (A) Experimental design. (B) Tumor growth curve of experiment described in A, n = 5 mice/group. (C) Experimental design and (D) tumor growth curve of experiment described in B, n = 5-8 mice/group. Data are representative of at least 2 independent experiments with similar results. ***P < 0.001 [Two-way ANOVA with Tukey correction].

### Reprogramming the tumor microenvironment with inactivated SARS-CoV-2 increases cross-presenting DCs and tumor antigen-specific CD8+ T cells

To determine the changes in the TIME following intratumoral administration of inactivated viruses in melanoma and breast cancer tumors, we performed flow cytometry analysis on treated tumor samples. Given that tumors with increased immune cell infiltration are associated with better response to immunotherapy and improved patient outcomes, we initially determined the presence of CD45^+^ cells within the tumor following treatment. In the B16 melanoma model, there was a significant increase in CD45^+^ cell infiltration from FluVx and iCOVID treatment. In the 4T1 breast cancer model, however, there was no observed difference in CD45^+^ cell infiltration amongst the three groups (Figure 2A & B). Because of the differences in immune cell infiltration between the two tumors models, we then assessed the changes in specific immune cell populations following each treatment. CD8^+^ T cell infiltration in the tumor has been strongly associated with improved tumor outcomes, thus we were particularly interested in the effects of intratumoral iCOVID on T cell infiltration especially when compared to FluVx. In melanoma and breast cancer models, intratumoral injection of iCOVID led to a significant increase in tumor infiltration of CD8^+^ T cells compared to controls, like that seen following influenza injection (Figure 2C & D). In addition to increased CD8^+^ T cell infiltration following iCOVID and FluVx administration, both treatments were observed to increase antigen presenting cell (APC) populations within the tumor. In the 4T1 tumor model, a significant increase in CD8^+^ cross-presenting dendritic cells was observed in both FluVx- and iCOVID-treated tumors compared to the control (Figure 3A)^7^. B16 melanoma tumors were found to have a significant increase in MHC-II^+^ cells in FluVx and iCOVID treated tumors (Figure 3B). These findings suggest that intratumoral iCOVID can shift the TIME in a manner that is similar to that of FluVx.

**Figure 2:**
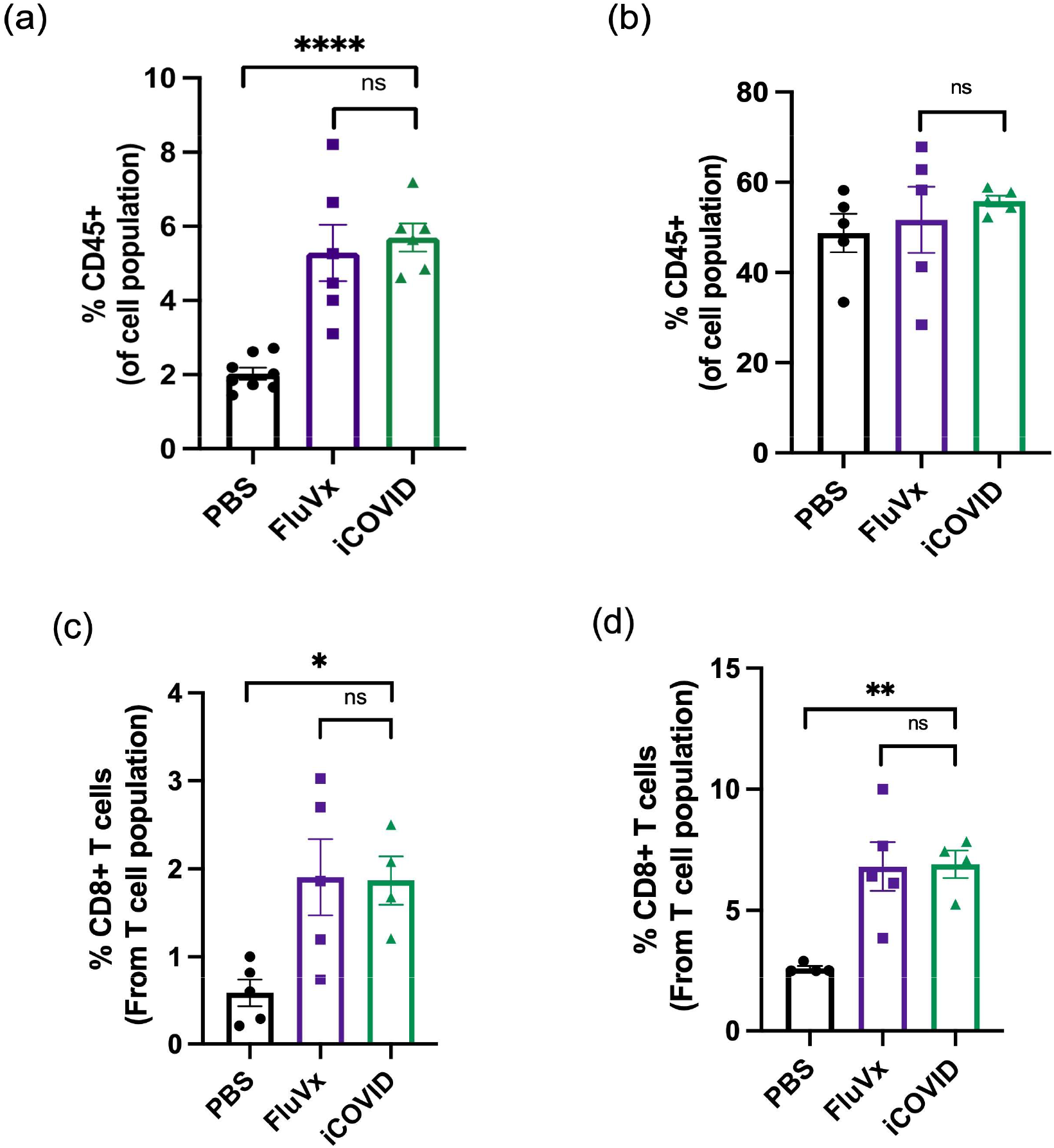
Intratumoral administration of inactivated SARS-CoV-2 increases the proportion of CD45^+^ and CD8^+^ T cells within the tumor microenvironment (A) in B16 melanoma tumor and (B) 4T1 TNBC tumor. (C) Intratumoral CD8^+^ T cells (CD45^+^CD3^+^) in melanoma tumor and (D) TNBC tumor. Data are representative of at least 2 independent experiments with similar results. ****P < 0.0001, **P < 0.01, *P < 0.05, ns P ≥ 0.05 [One-way ANOVA with Tukey correction].

**Figure 3:**
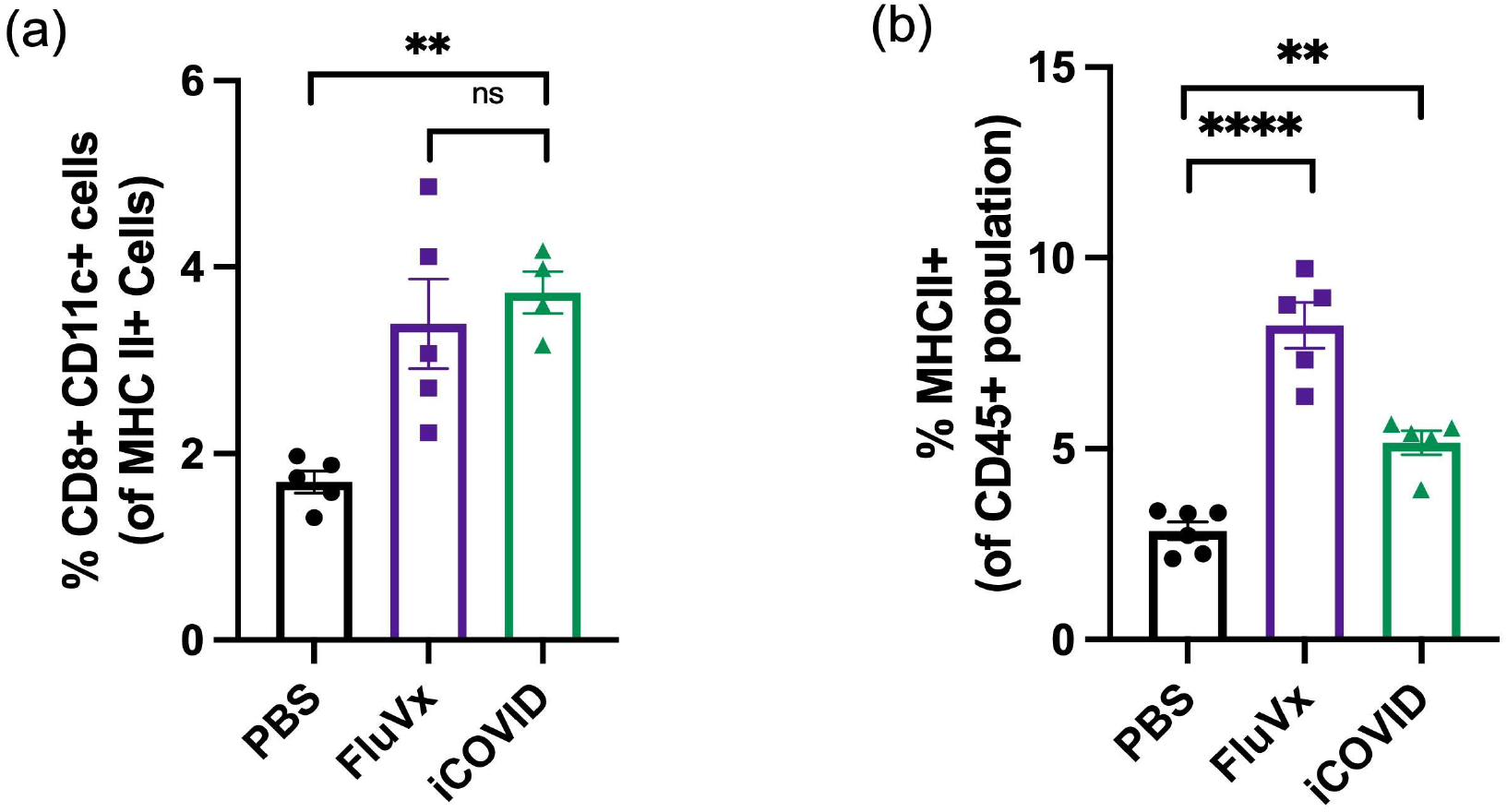
Intratumoral administration of inactivated SARS-CoV-2 increases intratumoral antigen presenting cell populations. (A) Cross-presenting dendritic cells (CD11c^+^ CD8a^+^) populations in 4T1 TNBC tumors after PBS, FluVx, or iCOVID treatment and (B) APCs (CD45+ MHC-II+) in B16 melanoma tumors after treatment. Data are representative of at least 2 independent experiments with similar results. **P < 0.01, ns P ≥ 0.05 [One-way ANOVA with Tukey correction].

### Inactivated SARS-CoV-2 treatment reduces Myeloid-Derived Suppressor Cell expansion in tumor microenvironment

Myeloid-derived suppressor cells (MDSCs) are known to suppress T cell responses in the setting of cancer, thus promoting an immunosuppressive tumor microenvironment^8^. Though not statistically significant, FluVx and iCOVID treatments both were observed to decrease CD11b^+^ Ly6G/Gr-1^+^ MDSC populations within 4T1 TNBC tumors (Figure 4A). A similar trend was observed in B16 melanoma tumors. Intratumoral FluVx and iCOVID administration both lead to a significant decrease in MDSCs in melanoma tumors (Figure 4B). These findings suggest that inactivated SARS-CoV-2 and influenza are both capable of relieving the tumor microenvironment of immunosuppressive MDSC populations.

**Figure 4:**
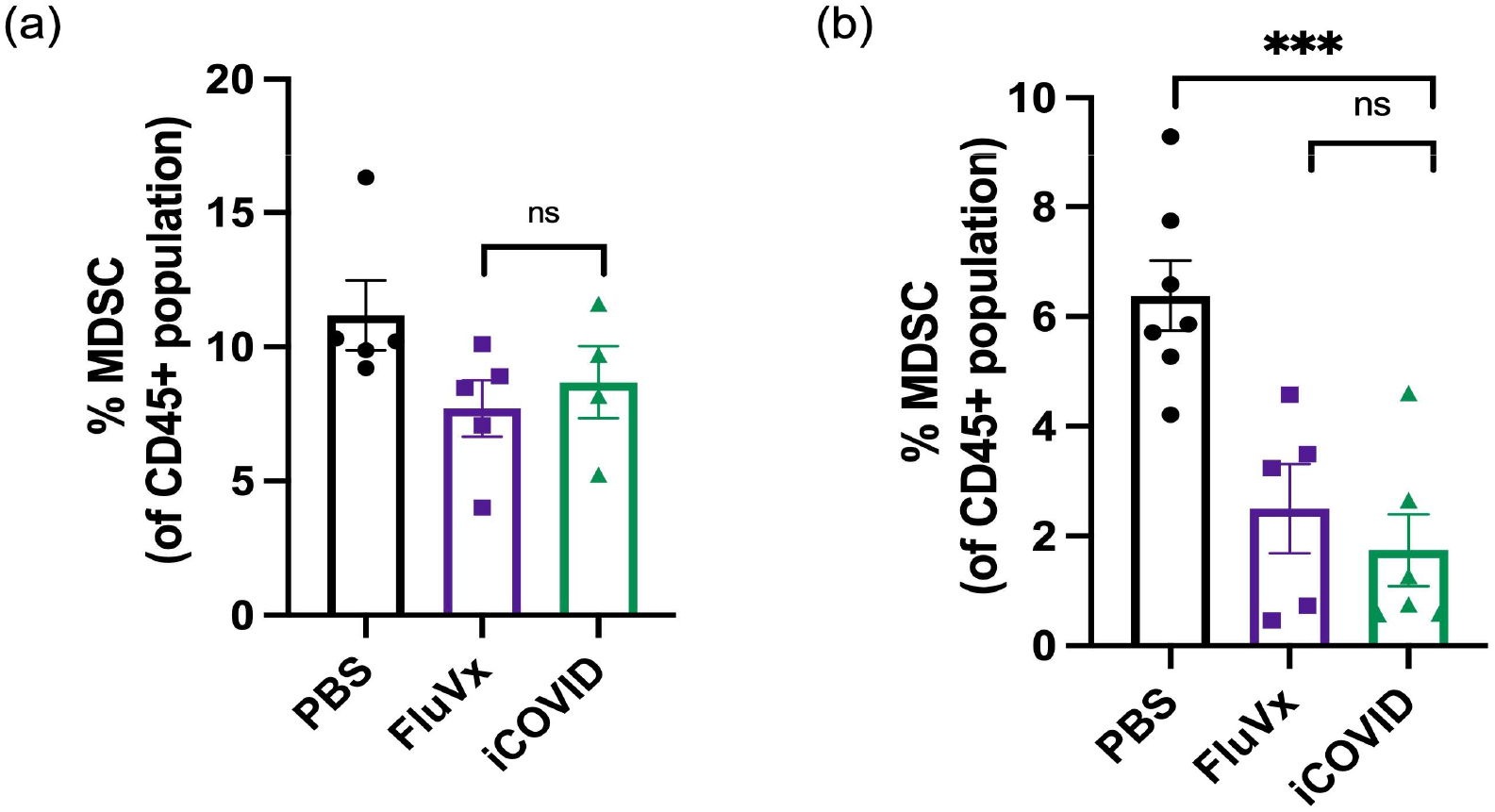
Intratumoral administration of inactivated SARS-CoV-2 decreases myeloid-derived suppressor cells (MDSCs) within tumor microenvironment. (A) Intratumoral MDSCs (CD11b^+^ Ly6G/Gr-1^+^) after FluVx, iCOVID, or PBS treatments in 4T1 TNBC tumors and (B) B16 melanoma tumors. Data are representative of at least 2 independent experiments with similar results. ***P < 0.001, ns P ≥ 0.05 [One-way ANOVA with Tukey correction].

### SARS-CoV-2-induced tumor regression is TLR7-dependent

TLRs identify microbial components, implying that TLRs function as adjuvant receptors to regulate both innate and adaptive immune responses^9^. It is possible that certain viral products act as TLR ligands and activate TLR7 signaling directly. Alternatively, like in the case of Drosophila host defense, viral infection may result in the production of endogenous ligands^10^. TLR7 activity was found to be increased in response to single-stranded RNA (ssRNA), a natural agonist found in the SARS-CoV-2 (Figure 5A), and the data were comparable to whicgh we discovered using FluVx^1^ (Figure 5B).

**Figure 5:**
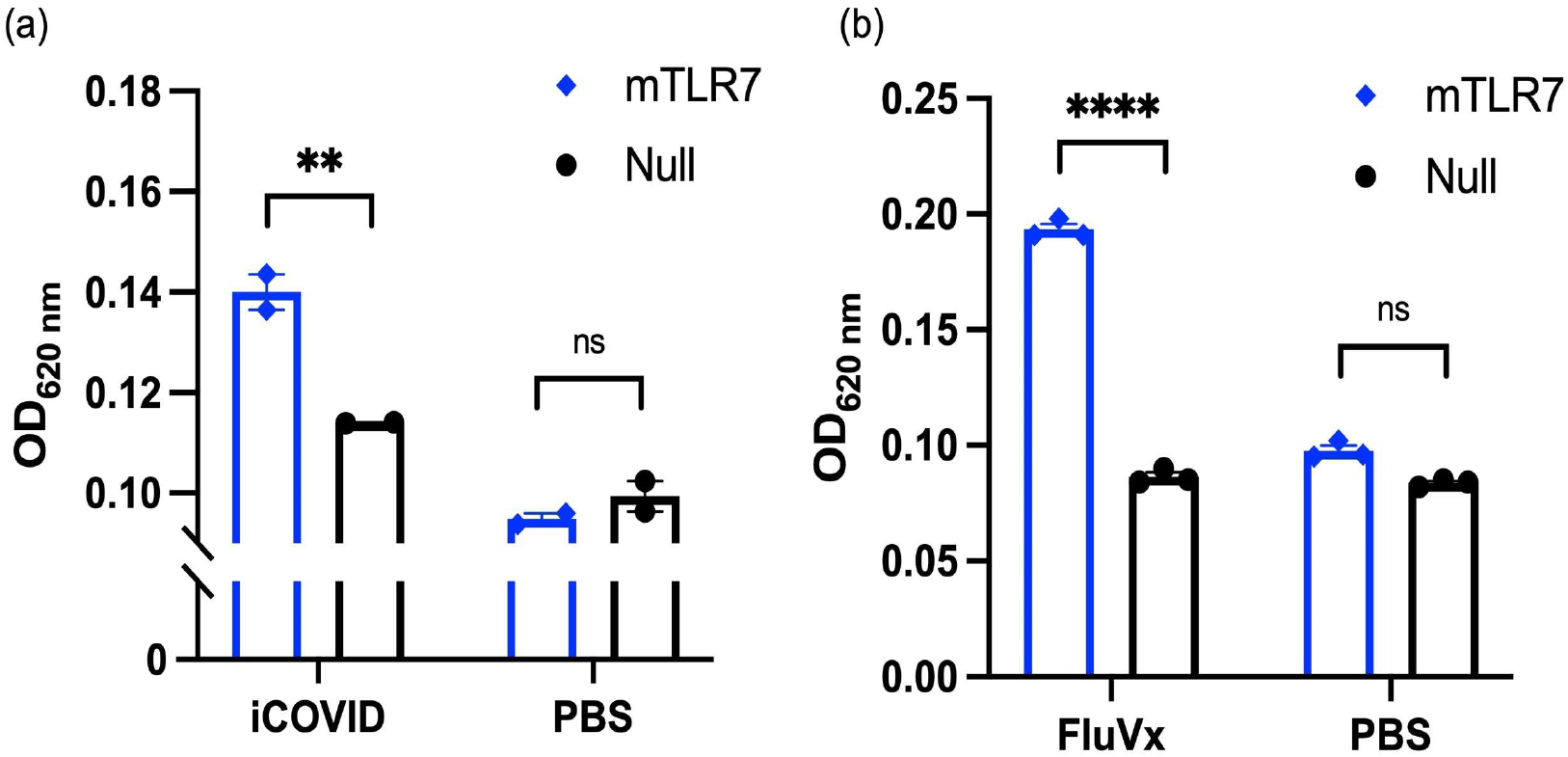
Inactivated SARS-CoV-2 increases TLR7 activity similar to inactivated influenza virus. (A) Bar graphs showing murine TLR7 stimulation following treatment with iCOVID or PBS. (B) Bar graphs are from one experiment run in duplicate wells and representative of at least two independent experiments with similar results. TLR7 activity was read at OD620 nm. ****P < 0.0001, **P < 0.01, ns P ≥ 0.05 [Two-way ANOVA with Sidak’s correction for multiple comparisons].

## Discussion

Clinical breakthroughs using immunotherapy to improve and prolong the lives of cancer patients have proven the immune system’s importance in cancer treatment. However, immunotherapies have only been able to induce long-lasting responses in a small percentage of patients thus far. As a result, new ways of engaging the immune system are required to take the next big step forward. Multiple lines of evidence suggest that local instigation of infectious microorganisms or synthetic molecules mimicking their components, the so-called pathogen-associated molecular patterns (PAMPs) acting on pattern recognition receptor agonists (PRRs), can be beneficial against cancers^11^, ranging from Coley’s clinical experiments with local treatment with bacteria or bacteria toxins to the generalized use of bacillus of Calmette and Guerin (BCG) for superficial bladder cancer^12^.

Multiple attempts have been made to target cytopathic effects against cancer utilizing so-called oncolytic viruses. A growing body of research suggests that replication-competent oncolytic viruses and replication-defective viral vectors work primarily by eliciting more effective antitumor immune responses. When the virus’s genome contains genes that boost immunity, such as cytokines and costimulatory factors, the virus’s potency can be increased. Intratumoral administration of viruses and virus-associated molecular patterns can result in anticancer effects, which are mostly mediated by the elicitation or potentiation of immune responses against the cancer.

According to recent findings in mice transplantable tumor models, the rotavirus, influenza, and yellow fever vaccinations are particularly well-suited to eliciting potent antitumor immunity against cancer after intratumoral injection^13^. Adenovirus, vaccinia virus, herpes simplex 1 virus (HSV), Newcastle disease virus, and reovirus intratumoral modified variations have showed exceptional activity in preclinical models and have moved, or are advancing, to the clinic^14^. T-VEC (Talimogene laherparepvec)^15^, an attenuated HSV engineered to encode granulocyte-macrophage colony-stimulating factor (GM-CSF), is approved for intratumoral infection in advanced melanoma and appears to have synergistic effects when combined with systemic treatment with anti-cytotoxic T-lymphocyte antigen 4 (CTLA-4) or anti-programmed death-ligand (PD-L1). In our research, we used SARS-CoV-2 to boost naturally weak antitumor immune responses, resulting in better cancer outcomes, in case of triple-negative breast cancer and skin cancer. Our findings show that inactivation of a non-oncolytic virus, such as SARS-CoV-2 can boost an anti-cancer immune response when supplied via intratumoral injection. This shows that the field of microbial-based cancer treatments (MBCTs), which has recently regained popularity, is not restricted to oncolytic pathogens or even the use of active pathogens^16, 17^. When homologous antigens are introduced within the tumor to induce an immune-”alarming” effect, virus-specific memory T cells have been proven to inhibit tumor growth in mice models^18^. Our findings suggest that injecting inactivated SARS-CoV-2 intratumorally reduces tumor growth by converting immunologically inactive cold tumors into immune-infiltrated hot tumors, thereby doubling the frequency of DCs (including cross-presenting DCs) and tumor antigen-specific CD8+ T cells in the tumor microenvironment.

